# Calorimetric analysis using DNA thermal stability to determine protein concentration

**DOI:** 10.1101/2023.09.25.559360

**Authors:** Matthew W. Eskew, Patrick W. Reardon, Albert S. Benight

**Affiliations:** ThermoCap Laboratories Inc, Portland, Oregon; Department of Chemistry, Portland State University, Portland, Oregon; OSU NMR Facility, Oregon State University, Corvallis, Oregon; Department of Physics, Portland State University, Portland, Oregon

## Abstract

It was recently reported for two globular proteins and a short DNA hairpin in NaCl buffer that values of the transition heat capacities, *Cp*,_*DNA*_ and *C*_*p*_,_*PRO*_ for equal concentrations (mg/mL) of DNA and proteins, are essentially equivalent (differ by less than 1%). Additional evidence for this equivalence is presented that reveals this phenomenon does not depend on DNA sequence, buffer salt, or T_m_. Sequences of two DNA hairpins were designed to confer a near 20°C difference in their T_m_’s. For the molecules, in NaCl and CsCl buffer the evaluated *C*_*p*_,_*PRO*_ and *C*_*p,DNA*_ were equivalent. Based on the equivalence of transition heat capacities, a calorimetric method was devised to determine protein concentrations in pure and complex solutions. The scheme uses direct comparisons between the thermodynamic stability of a short DNA hairpin standard of known concentration, and thermodynamic stability of protein solutions of unknown concentrations. In all cases, evaluated protein concentrations determined from the DNA standard curve agreed with the UV-Vis concentration for monomeric proteins. For samples of multimeric proteins, streptavidin (tetramer), Herpes Simplex Virus glycoprotein D (trimer/dimer), and a 16 base pair DNA duplex (dimer), evaluated concentrations were greater than determined by UV-Vis by factors of 3.94, 2.65, and 2.15, respectively.

## INTRODUCTION

Classical techniques for determining concentrations of DNA and protein solutions include absorbance spectroscopy (UV-Vis), the BCA, Bradford, Lowry and fluorescence assays which have the added requirement of specific molecular probes [1-3]. UV-Vis, either from direct measurements or a colorimetric assay, is the most widely employed method for quantitative determination of concentrations of nucleic acids and proteins in solution. To determine accurate protein concentrations for a specific molecule, these methods typically require pure solutions. Even though standard methods can generally provide reliable measurements of protein concentrations, they can yield conflicting results [4]. For many cases UV-Vis measurements can provide accurate concentrations. However, for complex solutions, such as those encountered in cell culture media and lysates, reliable concentrations from UV-Vis measurements can be tenuous and generally only provide a semi-quantitative estimate of the total protein concentration.

It has been well established that thermodynamic analysis of the temperature induced melting transitions of proteins and DNA provides a novel means to explore many aspects of molecular structural composition [5-13]. Calorimetric measurements of the transition provide a direct reflection of the collective forces responsible for molecular stability [6, 11, 14-16]. Many of the features related to molecular processing in biology such as ligand binding and intermolecular interactions can be probed, with unique information extracted, using calorimetric measurements [16-18].

Thermal denaturation or melting experiments are performed in a differential scanning calorimeter (DSC). In DSC melting experiments for DNA or proteins the heat capacity, C_p_, at constant pressure is measured as a function of temperature. A plot of C_p_ versus T is the DSC melting curve or thermogram. For DNA and many proteins, the heat induced melting transitions are endothermic with a pseudo-Gaussian shape. The temperature at the maximum peak height is defined as the transition temperature, T_m_. Likewise, the value of C_p_ at the maximum peak height is the transition heat capacity.

Here we report on the use of melting calorimetry to determine unknown concentrations of protein solutions. It is founded on the recent discovery for equal mass concentrations (mg/mL) of DNA or protein that their heat capacities, determined from the DSC thermogram peak heights at the transition temperatures, are the same i.e. *C*_*p*_,_*PRO*_ = *C*_*p,DNA*_ [19]. Based on this relationship a novel scheme was developed that enabled quantitatively accurate determinations of targeted protein concentrations in both pure solutions and impure complex mixtures.

Readily available synthetic short DNA oligomers with virtually any sequence provide the opportunity to fine-tune, by design, DNA secondary structure and corresponding sequence dependent thermodynamic stability. For example, several sequence dependent features of the thermodynamic stability of short DNA intramolecular hairpins have been well characterized [7, 20, 21]. As shown here, a DNA hairpin was chosen to serve as the stability standard for comparisons.

For the specifically designed and well characterized intramolecular hairpin that was used, DSC thermograms were measured for the DNA as a function of concentration over a 100-fold range. Plotting the thermogram peak heights versus DNA concentration produced a standard curve. By comparing the peak height on the thermogram for a protein at unknown concentration to the standard curve, and accounting for differences between the partial specific volumes of proteins and DNA, it is possible to directly evaluate the concentration of the protein.

## MATERIALS AND METHODS

### Reagents

Buffer solutions contained either NaCl or CsCl at 150 mM in 10 mM potassium phosphate, 15 mM sodium citrate adjusted to pH = 7.4 with HCl. Pure Proteins: Human Serum Albumin (HSA) (≥ 99% pure, Lot number: SLBT8667), Lysozyme (recombinant, expressed in rice, Lot number: SLCH2681), and Plasma (Human, Lot number: SLBT0202) were purchased from Sigma Aldrich (St. Louis, MO) and received as lyophilized power. Lysate from *E. coli* was purchased from Bio-Rad (Hercules, CA) and received as a lyophilized powder. The above proteins were prepared in buffer and stored at 4°C for at least 24 hours before use. Protein concentrations were confirmed by UV-Vis at A_280_ [22, 23].

Samples purchased from RayBiotech (Peachtree Corners, GA) included: HEK293 cellular supernatant, Lot number: 03U27020D. Unpurified angiotensin converting enzyme 2 (ACE2) expressed in human embryonic kidney cells (HEK293) received in the preparation media Lot number: 02U2802LL. Isolated, purified ACE2 Lot number: 04U27020GC. Purified receptor binding domain (RBD) from SARS-CoV-2, Lot Number: 05U22020TWB. Purified Human Herpes Simplex Virus glycoprotein D (HSV-GpD), Lot Number: 01U25022AG. All HEK293 solutions had a volume of 200 μL.

### DNA

Two DNA hairpins were designed to have T_m_’s that differed by approximately 20 °C. The “high-temp” hairpin was formed from the 20-base sequence 5’-CGG GCG CGT TTT CGC GCC CG-3’. The “low-temp” hairpin also had 20-bases with the sequence 5’-CGA TCG CGT TTT CGC GAT CG-3’. DNA strands were purchased from IDT (Coralville, IA) and received following their standard desalting routine. Lyophilized DNA was resuspended in either buffered NaCl or CsCl solutions and stored at 4°C. DNA concentrations were determined spectrophotometrically at A_260_ and agreed with vendor specifications. Figurative structures and sequences for the high- and low-temp hairpins are depicted in Figure 1(a, b).

**Figure 1:**
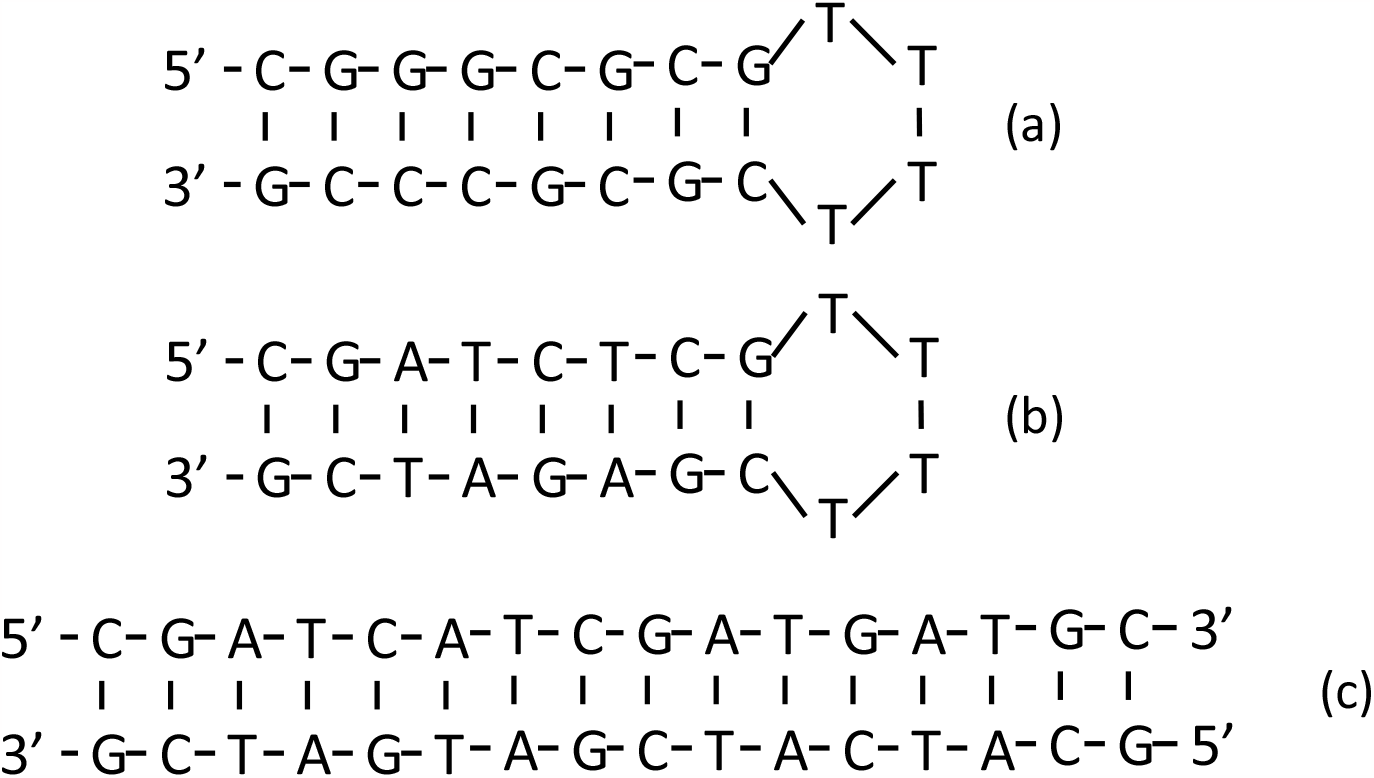
DNA Structures: (a) high-temp hairpin, (b) low-temp hairpin, and (c) duplex DNA

The 16 base-pair duplex DNA was formed from two individual stands whose base sequences are complementary. Duplex strands were purchased from IDT (Coralville, IA). The 5’ sequence of the duplex is 5’-CGA TCA TCG ATG ATG C-3’. Following their standard desalting routine, the preformed duplex was received as a lyophilized powder. The double-stranded DNA was resuspended in the desired appropriate buffer and stored at 4°C. DNA concentration was determined by UV-Vis at A_260_. The duplex is depicted in Figure 1(c).

### Calorimetric Measurements

Melting experiments were performed on a CSC differential scanning microcalorimeter (now T.A. instruments, New Castle, DE). Samples were prepared by adding specific components to buffer or pre-prepared media solutions in a sample volume of 0.5 mL. For DSC melting experiments, the sample heating rate was approximately 1 °C/min while monitoring changes in the excess heat (microcalories) of the sample versus temperature [16, 24, 25]. In their primary form, melting curves or thermograms of protein and DNA samples were displayed as plots of changes in microcalories versus temperature. For complex media experiments all DSC measurements were made on samples residing in the lysate media, either *E. coli* lysate or HEK923 supernatant. DSC thermograms were measured as a function of concentration for the high and low-temp hairpins, HSA and lysozyme. For each experiment, the transition temperature, T_m_, was determined by the temperature of the maximum peak height, 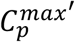on the thermogram.

### Data Reduction and Analysis

In primary form thermograms of proteins are displayed as plots of microcalories (*μcal*) versus temperature. Baselines of all thermograms were treated in an identical manner. Initially, for all mixtures the buffer baseline was first subtracted from the raw curves. Then baselines of the raw thermograms of *ΔC_p_* or *μcal* versus temperature were determined using a four-point polynomial fit, over the temperature range of the transition. The result of the analysis produced baseline corrected thermograms used for comparisons and analysis [16, 17, 19]. For all thermograms of proteins in complex media, the thermogram for media alone served as the background that was then subtracted from thermograms of samples containing the target protein in media [18]. These difference thermograms were then used to evaluate the concentration of desired protein components in the mixture.

### Statistical Analyses

For nearly all samples thermograms were collected at least in duplicate. Reproducibility and accuracy were verified using an ANOVA p-test. Statistical analysis and curve fitting were performed using Origin (Pro) Software, Version 2021b (Origin Corporation, Northhampton, MA). Linear fits of plots were quantified using a Pearson test [26, 27].

### Partial Specific Volumes of DNA

PSV values for the DNA hairpins and duplex DNA were determined by Analytical Ultracentrifugation (AUC). All sedimentation equilibrium experiments were performed in a Beckman Coulter Optima XL-A analytical ultracentrifuge (Beckman Coulter, Brea, CA) at 20°C equipped with absorbance optics. Data were measured at 260 nm in 6-channel cells. Samples were centrifuged at 24,000, 32,000, and 40,000 rpm, with a minimum equilibration time of 30 hours per speed. Buoyant molecular weights were determined using Heteroanalysis and PSV values were calculated using the Svedberg equation [28]. All experiments were conducted in standard NaCl buffer, with a density of 1.005584 g/ml.

## RESULTS

### The Transition Heat Capacity, *C*_*p*_,_*x*_

Under appropriate experimental conditions and standard data analysis procedures, the maximum peak height on measured thermograms of *ΔC_p_*versus temperature, 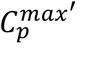 for *x* = DNA or proteins (PRO) can be given by [19, 29],

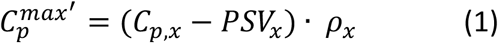

Where *ρ*_*x*_ is the mass concentration, *C*_*p,x*_ and *PSV*_*x*_ are the *transition heat capacity* and partial specific volume, respectively. It should be noted that *ρ*_*x*_ is the mass concentration of the sample reported as mg/mL. For the DNA and protein samples examined, an identical procedure was followed to evaluate *C*_*p,x*_ . Eqn (1) indicates a linear plot of 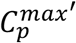versus concentration, *ρ*_*x*_, has a slope of *C*_*p,x*_− *PSV*_*x*_ [19]. For these samples thermograms were measured as a function of DNA concentration over the approximately 100-fold range from 0.02 to 2.5 mg/mL. All molecules were melted in buffer containing NaCl. To evaluate whether buffer composition played a role in the equivalence of transition heat capacity, some were also melted in buffer with CsCl replacing NaCl. For every sample, plots of 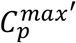 versus *ρ* were constructed. From linear fits of these plots, *C*_*p,x*_values were determined. Samples that were analyzed are listed in Table 1 along with results of the linear fits. As seen in Table 1, R^2^ values (Pearson test) were greater than 0.99 in all cases, indicative of highly accurate linear fits. Note, these fits were not forced to pass through the origin. This turned out to be of little consequence since values reported in Table 1 were statistically equivalent for fits forced through zero. These results are shown in Table S1 of the Supplemental Material.

**Table 1:** Linear Fits.

| Table 1: Linear Fits |  |  |  |  |
| --- | --- | --- | --- | --- |
| Sample | PSV (mL/g) | Slope ($\mu\text{cal mL/mg}$) | Intercept | $R^2$ |
| High Temp DNA Hairpin (NaCl) | 0.557 | $2.310 \pm 0.033$ | $0.119 \pm 0.044$ | 0.9988 |
| High Temp DNA Hairpin (CsCl) | 0.557 | $2.337 \pm 0.057$ | $0.040 \pm 0.096$ | 0.9982 |
| Low Temp DNA Hairpin (NaCl) | 0.557 | $2.322 \pm 0.005$ | $0.070 \pm 0.010$ | 0.9999 |
| Low Temp DNA Hairpin (CsCl) | 0.557 | $2.348 \pm 0.095$ | $0.050 \pm 0.173$ | 0.9951 |
| Duplex DNA (NaCl) | 0.625 | $4.140 \pm 0.024$ | $-0.082 \pm 0.041$ | 0.9999 |
| Human Serum Albumin (NaCl) | 0.733 | $2.122 \pm 0.032$ | $-0.098 \pm 0.033$ | 0.9996 |
| Human Serum Albumin (CsCl) | 0.733 | $2.248 \pm 0.098$ | $-0.156 \pm 0.129$ | 0.9962 |
| Lysozyme (NaCl) | 0.703 | $2.115 \pm 0.104$ | $-0.256 \pm 0.108$ | 0.9951 |

From the slopes given in Table 1 values of *C*_*p,x*_were determined for the DNA and protein samples. These are listed in Table 2. Previously, similar results were reported for the high-temp hairpin, HSA and lysozyme [19]. For the present study, new data was collected for these molecules. Where repeat experiments were conducted (high-temp hairpin, HSA and lysozyme in NaCl buffer) results were identical to those previously reported [18].

**Table 2:** Transition heat capacity *C*_*p,x*_ for DNA and proteins in standard (NaCl) and cesium chloride (CsCl) buffers.

| Molecule | $T_m$ ($^{\circ}\text{C}$) | $C_{p,x}$ ($\text{mcal} \cdot \text{g}^{-1} \cdot \text{K}^{-1}$) |
| --- | --- | --- |
| High Temp DNA Hairpin (NaCl) | $94.44 \pm 0.09$ | $2.86 \pm 0.03$ |
| High Temp DNA Hairpin (CsCl) | $94.36 \pm 0.06$ | $2.88 \pm 0.06$ |
| Low Temp DNA Hairpin (NaCl) | $75.72 \pm 0.13$ | $2.87 \pm 0.01$ |
| Low Temp DNA Hairpin (CsCl) | $75.56 \pm 0.06$ | $2.89 \pm 0.09$ |
| Duplex DNA (NaCl) | $71.55 \pm 0.07$ | $5.50 \pm 0.05$ |
| Human Serum Albumin (NaCl) | $63.85 \pm 0.14$ | $2.86 \pm 0.03$ |
| Human Serum Albumin (CsCl) | $63.16 \pm 0.13$ | $2.89 \pm 0.09$ |
| Lysozyme (NaCl) | $64.77 \pm 0.08$ | $2.82 \pm 0.13$ |
| Streptavidin (NaCl) | $77.39 \pm 0.05$ | 9.19 |
| HSV – glycoprotein D (NaCl) | $57.46 \pm 0.02$ | 6.33 |

As depicted in Fig 1, both 20 base DNA strands form intramolecular hairpins with an 8 base-pair stem with a T_4_ loop. For this study, the DNA duplex region of the original high-temp hairpin was re-designed to be relatively less stable by substituting A-T base pairs for G-C base pairs in the original sequence. The new sequence conferred a much lower Tm = 75.72 °C for the low-temp hairpin, compared to 94.44 °C. Thermograms measured as a function of concentration for the high-temp hairpin are shown Fig 2a. Similar plots for the other samples are given in the Supplemental Material. Note, for each hairpin the T_m_ was identical at all concentrations, behavior that is consistent with melting of a unimolecular tertiary structure. As shown in Fig 2a, peak heights, 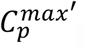, for the high-temp hairpin exhibited a linear response to DNA concentration. This is clear in Fig 2b where 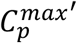 values for the high-temp DNA hairpin are plotted versus concentration.

**Figure 2:**
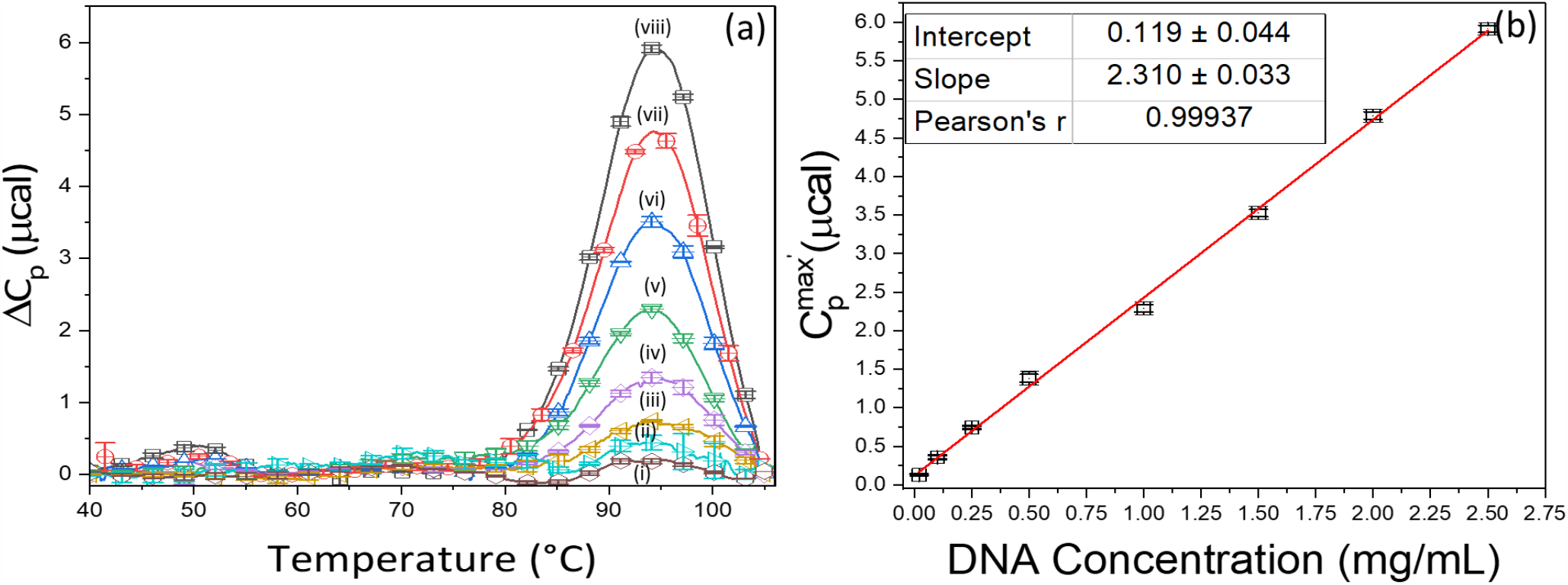
(a) Average thermograms for high-temp DNA in mg/mL: (i) 0.02, (ii) 0.10, (iii) 0.25, (iv) 0.50, (v) 1.00, (vi) 1.50, (vii) 2.00, (viii) 2.50. (b) Transition heat capacity curve for high-temp DNA.

From the slope of these linear plots (Table 1) and PSV for DNA it is possible to determine the transition heat capacity *Cp,_DNA_*. Reported PSV values vary for DNA from 0.57 ± 0.03 mL/g with the standard value assumed to be 0.55 mL/g [30, 31]. From AUC measurements for the DNA hairpins, PSV_DNA_= 0.557 ± 0.025 mL/g. With this value the transition heat capacity *Cp,_DNA_*= 2.86 ± 0.07 mcal g^-1^ K^-1^. Thermograms and linear plots like those shown Fig 2 were also obtained for the low-temp hairpin and HSA (Figs S1 and S2, Supplemental Material). The *Cp,_DNA_*values for these samples were in precise agreement with the value for the high-temp hairpin found here, and reported earlier for different samples of the same DNA and HSA [19]. Despite differences in their sequences, and corresponding different thermodynamic stabilities, *C*_*p*_,_*DNA*_values for the DNA hairpins were equivalent and independent of both sequence and T_m_ and equal to *C*_*p*_,_*PRO*_ for HSA at the same concentration [19].

### NaCl versus CsCl

Effects of electrolyte size and/or mass on the values of *C*_*p*_,_*DNA*_ and *C*_*p*_,_*PRO*_ were investigated by performing analogous experiments with CsCl replacing NaCl in the melting buffer. For these experiments, stock solutions of the high- and low-temp DNA hairpins and HSA were also prepared in CsCl buffer. In the same manner as in NaCl buffer, thermograms were measured as a function of DNA concentration from 0.5 to 2.5 mg/mL. Then for the molecules in CsCl, plots of 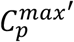 versus concentration were constructed (as shown in Fig 2b for the high-temp hairpin in NaCl buffer).

Slopes of plots of 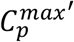 versus concentration yielded essentially identical *C*_*p*_,_*DNA*_values for the high-temp (2.88 ± 0.06 mcal g^-1^ K^-1^) and low-temp (2.89 ± 0.09 mcal g^-1^ K^-1^) hairpins in CsCl buffer. As displayed in Table 2, *C*_*p*_,_*DNA*_and *C*_*p*_,_*PRO*_ values were in precise agreement with those measured for the DNA hairpins and for the HSA sample in CsCl, *C*_*p*_,_*PRO*_ = 2.89 ± 0.09 mcal g^-1^ K^-1^. These values of *C*_*p*_,_*DNA*_and *C*_*p*_,_*PRO*_ determined in buffered NaCl or CsCl were identical. It should be noted in Table 2 for Streptavidin, HSV-GpD, and duplex DNA, evaluated transition heat capacities, *C*_*p*_,_*PRO*_ and *C*_*p*_,_*DNA*_were 9.19, 6.33 and 5.50 mcal·g^-1^·K^-1^, factors of 3.19, 2.2 and 1.91 larger, respectively, than the average over the other molecules. For streptavidin and HSV-GpD samples 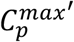 values measured at a single concentration were used to generate a full curve of 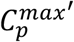 versus concentration by linear interpolation through the origin. From the slope of these plots *C*_*p*_,_*PRO*_ in Table 2, was evaluated. As posited below, the apparent discrepancies for streptavidin, HSV-GpD and duplex DNA were likely due to their multimeric composition.

### Determination of Unknown Protein Concentrations

With *C*_*p,DNA*_= *C*_*p*_,_*PRO*_, a scheme was devised to determine unknown concentrations of proteins in both pure and complex media solutions. Central to this process is construction of the standard curve. In the present case, from the collection of plots of 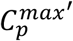 versus concentration that were constructed for the DNA hairpins and HSA, the plot for the high-temp hairpin was chosen as the standard curve. This choice is somewhat arbitrary. Analogous plots for the other molecules could have served equally as well. Once the standard curve was constructed, the sole input required to determine the unknown concentration of a protein is the peak height from a single thermogram for the target protein. The unknown concentration of the protein is the point on the ordinate axis on the DNA standard curve corresponding to 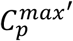 of the protein on the abscissa of the standard curve.

Accurate evaluations of protein concentrations, by comparison of diagnostic results for protein solutions of unknown concentrations with the DNA standard curve, also requires a correction that accounts for differences between PSV values for DNA and proteins. While the transition heat capacities for HSA and lysozyme, *C*_*p*_,_*PRO*_ and DNA, *C*_*p,DNA*_ in Table 2 are essentially equivalent, the average PSV for proteins are different. As measured by AUC, the PSV_DNA_= 0.557 ± 0.025 mL/g while most globular proteins have PSV values that vary from 0.70-0.75 mL/mg [32-36]. Also, for independent lots of duplex DNA the evaluated PSV was 0.623 ± 0.004 mL/g, considerably larger than what has been reported for duplex DNA [30]. In fact, differences in the slopes of plots of 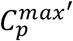(Protein) or 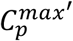 (DNA) versus concentration (Fig 2b, Table 1) are due directly to differences between PSV values for proteins and DNA.

After accounting for differences in PSV values for DNA and protein, the protein concentration is obtained. As will be shown below this process provides semi-quantitative estimates of protein concentrations in both pure and complex media.

From rearrangement of Eqn (1) the *transition heat capacity* is given by,

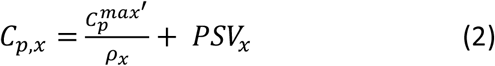

The analytical process is founded on the equality of values of the transition heat capacities for protein and DNA, determined by Eqn (2). Since *C*_*p*_,_*PRO*_ = *C*_*p,DNA*_

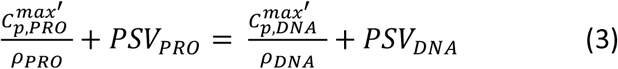

Solving for the unknown protein concentration,

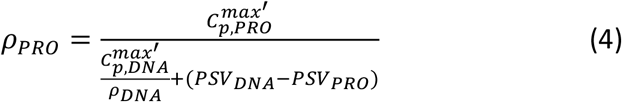

Eqn (4) reveals protein concentration, *ρ*_*PRO*_, can be evaluated from a single 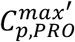 for a target protein. Practically, reasonable estimates of protein concentration can be obtained from a single DNA thermogram of known concentration. Provided that 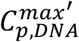 is taken from DNA thermograms measured at concentrations above 1.0 mg/mL since larger deviations occur at smaller DNA concentrations.

More accurate concentration determinations can be made using the complete DNA standard curve. The standard curve displayed in Fig 2b was constructed from multiple measurements of thermograms for the DNA standard as a function of concentration. The slope of the best fit line to the standard curve, 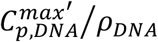, was evaluated and inserted in Eqn (4). Then after accounting for the difference between the partial specific volumes for the DNA and protein, *PSV*_*DNA*_−*PSV*_*PRO*_, concentration of the protein *ρ*_*PRO*_, was determined from 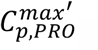 .

Eqn (4) was employed to determine the protein concentrations in several solution contexts. In what follows, examples include proteins in pure solutions, plasma and complex fluids such as cellular media. Sample preparation and experimental and analytical protocols were as described in Materials and Methods.

### Pure Solutions

The DNA standard curve displayed in Fig 2b was used to determine mass concentrations for several proteins and a few complex fluids. Some of which were also examined previously [18]. PSV values given in Table 3 for HSA [16, 19, 22], lysozyme [2,4], SARS-CoV-2 RBD [37-39], ACE2 [37], streptavidin [40], and HSV-GpD [41] were taken from the literature. Table 3 shows for all cases examined, there was excellent agreement between protein concentrations estimated from the DNA standard curve (column 3) and concentrations determined by UV absorbance or measured dry weight (column 4). Instead of an average over all peak heights at a particular concentration, peak heights in Table 3 were randomly selected from a set of peak heights obtained from the ranges of concentrations measured for each sample.

**Table 3:** Protein Concentration Comparison.

| Sample | Peak Height (μcal) | PSV (mL/g) | DNA Assay (mg/mL) | Independent Concentration (mg/mL) |
| --- | --- | --- | --- | --- |
| HSA | 3.70 | 0.733 | 1.74 | 1.79 |
| Lysozyme | 3.55 | 0.703 | 1.65 | 1.80 |
| SARS-CoV-2 RBD | 0.75 | 0.71 | 0.35 | 0.40 |
| ACE2 | 0.49 | 0.71 | 0.23 | 0.20 |
| HSA in Plasma | 2.08 | 0.733 | 0.98 | 1.08 |
| *E. coli* Lysate | 3.89 | 0.725 | 1.82 | 2.00 |
| ACE2 unpurified | (C)0.09,<br>(N) 0.06 | 0.69 | (C) 0.04,<br>(N) 0.03 | ----- |
| Streptavidin | 6.71 | 0.70 | 3.11 (3.94x) | 0.79 |
| HSV-GPD | 2.23 | 0.74 | 1.06 (2.65x) | 0.40 |
| 16mer dsDNA | 4.09 | 0.623 | 1.83 (2.15x) | 0.85 |

Overall, resulting concentrations evaluated from DSC measurements of proteins and the DNA standard curve differed (on average) by less than 8% from those determined by UV-Vis or dry weight.

In a prior study, two additional pure proteins, SARS-CoV-2 receptor binding domain (RBD) and angiotensin converting enzyme 2 (ACE2) were also examined [37-39]. Evaluated concentrations for these proteins agreed with independently determined reported values. Examination of Table 3 reveals for most samples, our method slightly underestimated the protein concentrations. The exception is ACE2 where the concentration was slightly overestimated by 13%. As previously noted, purified ACE2 samples displayed a single peak on their thermograms, in contrast to the expected two-peak behavior exhibited by native ACE2 [42]. Regardless, Table 3 indicates good agreement for concentration determinations for a variety of proteins in pure and complex media solutions, and DNA. Exceptions were observed for Streptavidin, HSV-GpD and short duplex DNA. As suggested below these exceptions are largely attributed to their higher molecular stoichiometries compared to the other molecules in Table 3.

Streptavidin is known to be a homo-tetramer comprised of four identical subunits (stoichiometry = 4) associated in a stable complex at 20°C regardless of concentration [43]. In Table 3 the concentration evaluated for streptavidin using the DNA assay (3.11 mg/mL) was also nearly a factor of four times (3.94) greater than the UV-Vis concentration of 0.79 mg/mL. To independently verify the streptavidin concentration, the commonly accepted extinction coefficient of 139,000 M^-1^cm^-1^ at 280nm was employed [43-45]. This value was used to determine the independent concentration of streptavidin given in column 4 of Table 3, also in agreement with the dry-weight mass used to prepare the protein solution. This observation is at odds with the calorimetric method (DNA assay) and apparently suggests the method evaluates molecular samples as monomers, regardless of their multimeric structure. This is the probable source of the discrepancy between the concentrations in Table 3 evaluated using the DNA assay and UV-Vis measurements. Alternatively, if the extinction coefficient for streptavidin monomers of 34,000 M^-1^cm^-1^ was used [43, 46] instead to evaluate the concentration by UV-Vis measurements, the same absorbance value would produce a concentration four times greater than using the tetramer extinction coefficient. However, since the mass concentration in the sample was the same, the four-fold lower molecular weight of the monomer must also be taken into account, which reduced the concentration by a factor of four, in agreement with our results. From these observations a significant benefit of the DNA assay emerges. It does not require prior knowledge of the sample to determine molecular stoichiometry.

It has also been reported that streptavidin is highly stable and even in the presence of SDS, temperatures above 60 °C are required to separate the tetramer into dimers and monomers [43]. Since SDS was not used here, even higher temperatures were required to dissociate the tetramer, consistent with other DSC experiments of streptavidin [47]. Once temperatures approach the T_m_, the tetramer dissociates to four monomers, increasing the actual monomer concentration by a factor of four. The four identical monomers each melt with a corresponding equivalent transition heat capacity. In this case the protein concentration evaluated from calorimetric measurements is approximately four times the tetramer concentration determined from absorbance measurements, and almost exactly what would be expected using the monomer extinction coefficient.

Another example was purified HSV-GpD. The evaluated concentration in column 4 of Table 3 was 1.06 mg/mL, a factor of 2.65 greater than the independently determined value of 0.40 mg/mL. Again, this finding is consistent with the known structure of HSV-GpD where there have been several reports that pure solutions of the protein contain both dimer (stoichiometry = 2) and trimer (stoichiometry = 3) forms [48-50]. In analogy to streptavidin, the ratio of 2.65 for HSV-GpD indicates that the protein solution was likely comprised of a mixture of dimer and trimer species, with slightly more trimer. This example provides additional support for the proposition that the DNA assay can also inform on the molecularity of multimeric proteins.

Likewise duplex DNA is a dimer comprised of two strands (stoichiometry = 2), whose base sequences are complementary. Concentrations determined by UV-Vis at 20°C assume a duplex (dimer) comprised of two monomer single strands. In analogy to streptavidin the duplex is assumed to be a “monomer” at 20°C, that then melts to two monomer single strands at higher temperatures. The duplex “monomer” concentration is twice the concentration of each single stand monomer. In this scenario the concentrations determined by UV-Vis at 20°C should be multiplied by the stoichiometry for duplex DNA.

As shown in Table 3 concentrations determined from the melting behavior of these molecules over the temperature range from 30 to 100°C, were 3.11 compared to 4 x 0.79= 3.16 mg/mL for streptavidin. For HSV-GpD containing a mixture of dimers and trimers compare 1.06 mg/mL to the range of 0.8-1.2 mg/mL. For duplex DNA compare 1.83 mg/mL to 2 x 0.85= 1.70 mg/mL. Results for these examples are all consistent with the notion that the higher *C*_*p*_,_*PRO*_ and *C*_*p,DNA*_ values, and higher concentrations determined from them, are indicative of higher stoichiometries of their tertiary structures, compared to the monomer. In effect, the DNA assay provides the monomeric concentration, regardless of oligomeric state. Thus, as demonstrated for the examples given above, the calorimetric method also provides insight into molecular tertiary structure where standard UV-Vis measurements cannot.

### Proteins in Complex Solutions

These experiments involved evaluating the concentration of a single protein target in complex solutions, as opposed to a single protein in buffer alone.

### Plasma

Human plasma served as an ideal test case. Over 9000 individual proteins comprise the plasma proteome, but a plasma thermogram is primarily due to seven major proteins [51, 52]. HSA comprises the major peak on a plasma thermogram. The height of this peak was used to estimate the fractional concentration of HSA in the complex mixture with other plasma protein components. As shown in Table 3 (row 5) there was good agreement between the reported fractional concentration of HSA in whole plasma 55-60% [22, 51-55], and the concentration of HSA extracted from the peak height on the whole plasma thermogram, estimated to be 0.98 mg/mL, or 54.4% of the total concentration of the plasma, 1.8 mg/mL. In addition to HSA, six other proteins can also contribute to the peak height in a minor way. It should be noted that because their individual thermograms significantly overlap, the HSA peak on the plasma thermogram centered at ∼62°C also contains slight contributions from the other six proteins most notedly alpha acid glycoprotein. However, since alpha acid glycoprotein (and the other proteins) comprise a small percentage (<1%) of the total concentration of human plasma, effects on the plasma thermogram compared to HSA are largely negligible [52].

### *E. coli* lysate

To mimic conditions comparable to those possibly encountered for the unpurified lysates of expression systems, lyophilized *E. coli* cellular lysate was purchased from Bio-Rad, and a known concentration was re-suspended in standard NaCl buffer. The thermogram for the *E. coli* lysate displayed a broad curve from ∼40-90°C with no significant distinguishing features [18]. The absolute height of this broad peak was used to estimate the concentration of lysate in the sample. Since the sample contains many unidentified proteins, a PSV value of 0.725 mL/g was used as the average of typical values reported for globular proteins (0.70-0.75 mL/mg) [32-36]. As Table 3 (row 6) shows, despite the complex milieu of the lysate, the peak height on the broad thermogram of the lysate alone, yielded a predicted concentration in agreement (within 10%) of the actual reported concentration of protein in the lysate solution. Although intriguing, this good agreement may be fortuitous and has not been independently substantiated.

### HEK293 supernatant

To explore the more realistic environment of an actual expression system, unpurified ACE2 was purchased and used for comparison with a purified ACE2 sample. Unpurified ACE2 contained ACE2 expressed from HEK293 cell lines in cellular supernatant. The thermograms measured for the supernatant alone and for supernatant containing an unknown amount of expressed ACE2 are displayed in Fig 3a. Both curves were broad over the temperature range displayed (40-100 °C), and the thermogram for ACE2 solution had a greater intensity for about 80% of the transition region. Subtracting the curves in Fig 3a provided the curve for ACE2 alone shown in Fig 3b. This curve clearly displayed two peaks around 55°C and 70°C. Also, shown in Fig 3b is the thermogram collected for purified ACE2 (200 μg/mL), that exhibits a single peak at ∼55°C [18, 42]. Subtracting background baseline from the extracted curve for AC2 shown in Fig 3b resulted in the curve for unpurified AC2 shown in Fig 3c, where the two peaks are clearly displayed. There is clear agreement with published results [42]. The thermogram in Fig 3c is very similar to the published thermogram for somatic bovine ACE2 [42].

**Figure 3:**
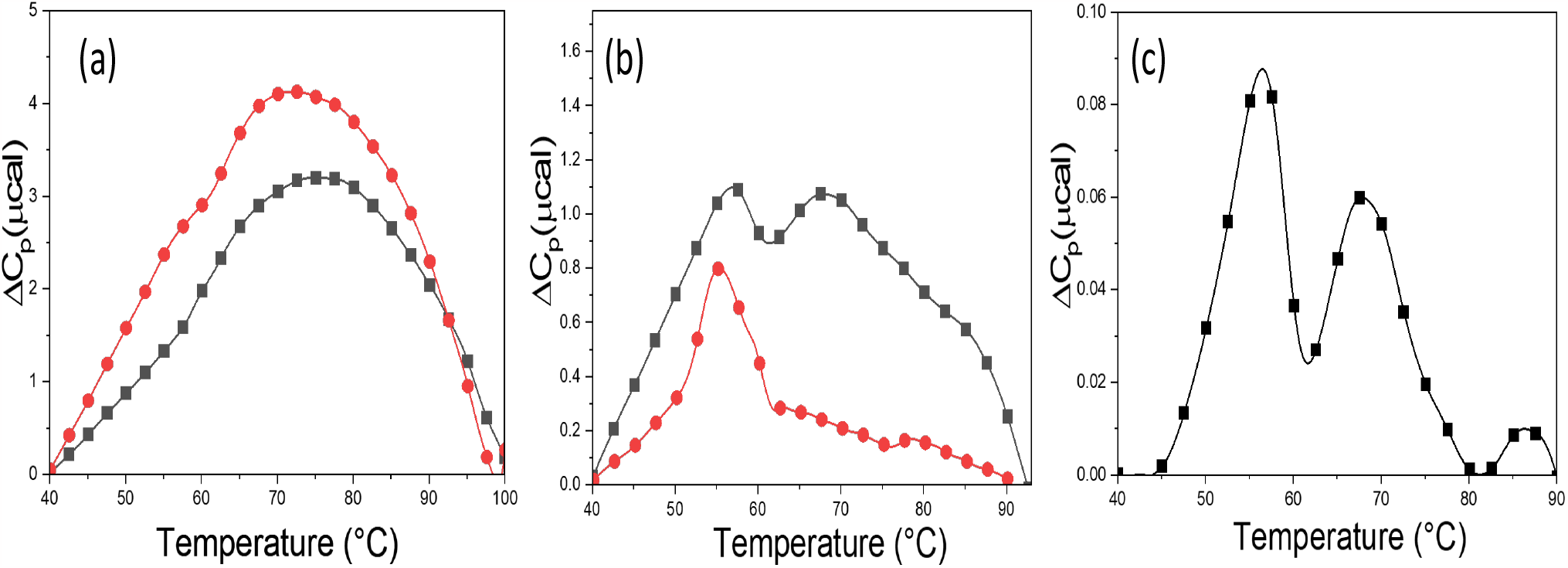
(a) Thermograms for the HEK293 cell supernatant background (black line) and a solution of the supernatant containing an unknown amount of expressed ACE2 (red line). (b) The difference thermogram corresponding to ACE2 protein alone obtained by subtracting the curves in (a) (black line); thermogram of a 0.20 mg/mL sample of purified ACE2 (red line). (c) Baseline corrected ACE2 curve from (b) and consistent with the reported thermogram for bovine somatic ACE2 [42].

Baseline subtracted thermograms were used to estimate the concentration contributions from each domain of the ACE2 sample in Table 3 (row 7). For the C-domain, the concentration corresponding to the peak at ∼55°C, was 0.04 mg/mL. Whereas for the N-domain, corresponding to the peak at ∼70°C, the concentration was 0.03 mg/mL. Our method was able to provide an estimate for the amount of ACE2 in the unpurified cellular supernatant of the HEK293 expression system. As it was unknown to them, the vendor was unable to provide the actual concentration of ACE2 in the unpurified sample. Accuracies of these concentration estimates await verification.

## DISCUSSION

### Equivalance of Transition Heat Capacities

Affirming evidence was obtained for the equivalence of the transition heat capacites at T_m_ for proteins, *C*_*p*_,_*PRO*_ and DNA, *C*_*p,DNA*_ at the same mass concentration. Two DNA hairpins with a nearly 20°C difference in T_m_ values were examined in NaCl buffer and had the same *C*_*p,DNA*_ Thermograms for these hairpins and HSA in CsCl buffer also provided identical results. It should be noted that the numerical values for the transition heat capacities, at the T_m_, shown in Table 3 differ from the reported heat capacity at 25°C for HSA, lysozyme and other globular proteins [5, 19, 29]. Based on the equivalence of *C*_*p*_,_*PRO*_ and *C*_*p,DNA*_, several novel applications have emerged.

### Concentration determination

A scheme was devised to determine unknown concentrations of individual proteins in pure and complex mixtures. Minimally, the scheme requires a single thermogram measured for both the protein of unknown concentration and DNA hairpin at a known concentration, along with the PSV values for DNA and proteins. Greater accuracy can be obtained if the entire DNA standard curve is employed, particularly at lower concentrations. The concentrations reported in Table 3 evaluated using the DNA standard curve agreed with concentrations determined by UV-Vis and dry weight measurements. It seems unlikely that consistent accuracy of evaluated concentrations, for such a wide variety of proteins, would have been possible if the transition heat capacities were not essentially equivalent. This was strictly true for all the monomer globular proteins examined (thus far). As discussed subsequently, additional evidence was provided suggesting the multimeric structures of streptavidin (a tetramer), HSV-GpD (dimer/trimer mix) and duplex DNA (a dimer) are also reflected in their transition heat capacities.

### Multimeric Proteins

Examples of monomeric proteins that were analyzed included HSA, Lysozyme, and SARS-CoV-2 RBD. The tertiary structures of these proteins form from a single polypeptide chain. Even though the tertiary structure of HSA contains three domains, the thermogram displays a single transition. Despite their ordered secondary and tertiary structures, the other proteins also displayed single peaks on their thermograms. In contrast, ACE2 is a heterodimer comprised of two different non-covalently associated monomeric subunits, the C- and N-domains [42, 56, 57]. These domains apparently differ enough in stability that the thermogram for ACE2 displayed two distinct peaks, presumably with each monomeric subunit displaying a distinguishable transition. For ACE2 the overall protein concentration in solution was expected to be 0.07 mg/mL. From the transition peak heights on the thermogram for ACE2 the evaluated concentrations were 0.04 mg/mL for the C-domain and 0.03 mg/mL of the N-domain for a total of 0.07 mg/mL for the total protein.

The values of the transition heat capacities found for streptavidin, HSV-GpD and linear duplex DNA were different from the other proteins and DNAs. Streptavidin is comprised of four identical protein monomers [44]. Unlike the other proteins that were examined, for streptavidin the evaluated concentration differed significantly from that determined from UV-Vis measurements. This concentration difference in Table 3 (row 8) was preserved over a range of concentrations and multiple experiments. Therefore, experimental error cannot account for the apparent discrepancy. The ratio of calorimetrically evaluated concentration to the UV-Vis determined concentration for the tetrameric protein was nearly fourfold higher (3.94) than observed for the monomeric proteins. Since all results were normalized by concentration, the observed fourfold difference in peak height was initially surprising but perhaps not unexpected considering the stoichiometric composition of streptavidin. In the case of streptavidin, the calorimetric method appears to be sensitive to not only contributions from separation of the four monomers in the overall secondary structure, but also melting of the four individual monomers comprising the tertiary structure. Likewise, for HSV-GpD the ratio of the evaluated concentration to the UV-Vis concentration was a factor of 2.65 suggesting a mixture of dimers and trimers at approximately, 35% and 65%, respectively.

A novel aspect of the calorimetric approach is the insights it can provide into tertiary structural composition of unknown proteins. This capability is not available with conventional spectroscopic methods such as UV-Vis and fluorescence. For example, as demonstrated here, calorimetric analysis was able to differentiate whether protein domains form from a contiguous primary structure i.e. monomeric (HSA, lysozyme); or are multimeric i.e. dimeric (unpurified ACE2), dimer and trimer mix (HSV-GpD) and tetrameric (streptavidin) formed by non-covalent association of identical monomer subunits. The calorimetric method evaluates the monomeric concentration, regardless of the multimeric state.

### Duplex DNA

Thermograms collected for the 16 base-pair double stranded DNA as function of concentration are shown in Fig 4. Note, the curves incrementally shift to higher temperature as the concentration increases. Behavior entirely expected for the concentration dependence of short linear duplex DNA melting [58, 59]. In precisely the same manner as for the hairpin DNAs and protein samples the DNA concentration was determined from the peak heights on DSC thermograms. The evaluated concentration was 2.15 greater than determined by UV-Vis. While one thermogram at a single concentration was used to evaluate the DNA concentration in Table 3, due to the linear relationship between the measured peak height and protein concentration, regardless of the concentration chosen, the resulting evaluated concentration was a factor of 2.15 greater than determined by UV-Vis. These results for short duplex DNAs are analogous to those found for streptavidin and HSV-GpD, and consistent with the bi-molecular composition of a DNA duplex dimer that forms via association of two independent single strand monomers. Although these are reasonable and compelling scenarios for streptavidin and HSV-GpD, further calorimetric examination of additional multimeric proteins and multi-stranded DNAs will be required before these interpretations can be generalized.

**Figure 4:**
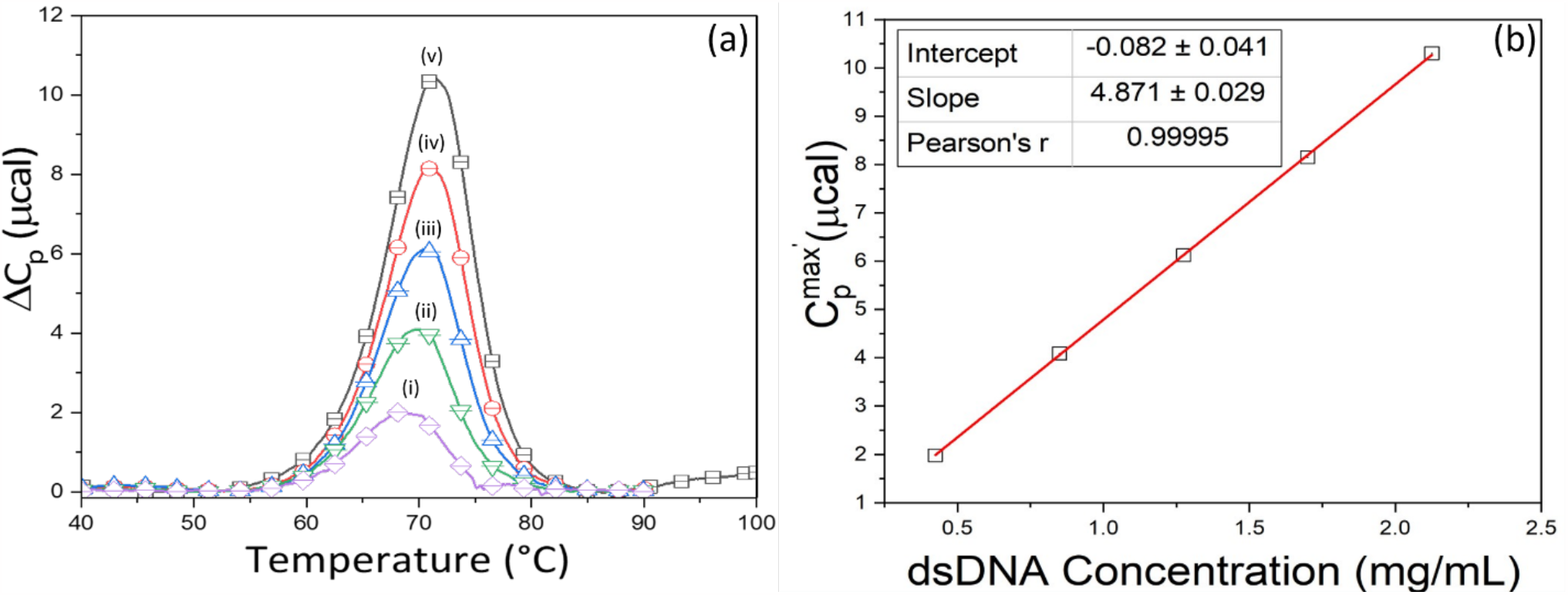
(a) Average thermograms for dsDNA concentrations in mg/mL: (i) 0.43, (ii) 0.85, (iii) 1.28, (iv) 1.70, (v) 2.13. (b) Transition heat capacity curve for dsDNA.

### A tool for Drug Discovery and Screening

Insight provided by the observed equivalence of the transition heat capacities for proteins and DNA provides for novel modern applications. For example, *in vitro* expression systems (prokaryotic and eukaryotic) are the source of many specially engineered low abundance molecules and biopharmaceuticals (biologics) [60]. Expression systems provide a means for generating important biological molecules (proteins, peptides, antibodies), on various scales, relatively fast for reasonable cost. Unfortunately, isolation and purification of expressed molecules can be expensive and time-consuming. For a moderately abundant and soluble protein, conventional purification strategies can require 27 or more individual steps and at least four days to complete [61]. In some cases, additional time-consuming purification, fractionation, and screening procedures may be required to reach desired levels of purity. This time requirement currently constitutes an obstacle to biologic drug development and emphasizes the need for a quick and quantitatively reliable means to reliably estimate unknown concentrations of protein products of *in vitro* over-expression systems. In principle, the demonstrated capability of calorimetric analysis could provide a means for quickly screening libraries of potential compounds for yield and activity, without requiring prior purification. In so doing, relieving a critical bottleneck in biopharmaceutical development.

Previously we demonstrated the utility of DSC to detect and analyze ligand binding to unpurified protein products of expression systems [18]. At that time, it was not possible to quantitatively evaluate ligand binding constants for proteins whose concentrations were unknown. As reported, this was the case for an unpurified protein (ACE 2) product of an expression system [18]. Now that we have demonstrated the ability to quantitatively evaluate the concentration of specific proteins in complex media mixtures, it should be possible to use DSC to quantitatively evaluate ligand binding constants to protein products of expression systems.

### DSC versus UV-Vis

The reported calorimetric DNA assay provides unique information that other methods such as absorbance measurements (UV-Vis) cannot. As has been demonstrated, the calorimetric scheme enables quantitative evaluation of unknown protein concentrations in pure solutions and complex mixtures, without the need for probes or binding assays. Insights into molecular stoichiometry of proteins and DNA can also be gleaned from calorimetric measurements. An added feature of calorimetric analysis is the much higher solution concentrations that can be analyzed compared to UV-Vis absorbance measurements.

UV-Vis measurements can be used to quantitatively evaluate concentrations of biological molecules. Practically, UV-Vis measurements require samples having sufficient optical activity in the UV, with measurable spectroscopic properties or the ability to bind a colorimetric probe. UV-Vis most commonly requires solutions, in a one cm pathlength cuvette, having UV absorbance ≤ 1 OD/ml. For more concentrated solutions, accurate absorbance measurements require dilutions to achieve readings ≤ 1 OD/ml. Additional sample handling involved in preparing diluted samples can also be an unnecessary source of error.

One practical difference and consequence of calorimetric analysis is that it requires melting of the protein sample. During the melting process, typically over the range from 30 to 90°C, the sample may get damaged, or irreversibly aggregate. Consequently, the sample may not be retrievable in active form after the experiment and consequently sacrificed. This is not the case for UV-Vis measurements at 20°C where the sample can usually be safely retrieved.

### Conclusions

Precise origins of the observed heat capacity equivalence for proteins and DNA remain a mystery, but none-the-less an empirical fact for the examples examined thus far. Several potential sources are merely mentioned here. The suspected role of hydration has been discussed and likely a prevalent component [14, 19, 32, 62-68]. In addition to hydration, electrostatic and vibrational effects can also contribute to the transition heat capacity [69, 70]. Our results in CsCl indicated no differences for the DNAs and HSA from what was found in standard NaCl buffer. Cesium has a much larger (50%) Van der Waals radius and 5.75 times greater mass (132.91 g/mol) than sodium (22.99 g/mol). If solvent interactions contributed significantly to *C*_*p,DNA*_ and *C*_*p,PRO*_ (then different values in NaCl versus CsCl might have been expected. None were observed suggesting electrostatics and associated polyelectrolyte effects on the observed equivalence must be relatively small. This is in accordance with earlier modeling studies suggesting electrostatics only contribute in a minor way to the heat capacity [69].

Several examples of our novel approach to calorimetric measurements and analysis have been previously reported [17-19]. As described here the DNA assay for evaluating concentrations of protein solutions from DSC thermograms is the most recent of these applications. Others have involved using calorimetric analysis to examine ligand binding reactions in both pure solutions and complex media mixtures [18]. The essence of our approach lies in the power of relative measurements which have been shown to provide a sensitive and accurate means to examine protein structure and ligand binding. For example, *ratiometric* analyses of DSC thermograms, versus absolute measurements, are much more accurate and informative, without the added requirement of tenuous model building, associated underlying assumptions and analysis [17-19].

Clearly the calorimetric approach provides unique capabilities with potentially powerful applications. With the novel information that can be obtained using calorimetry important applications in drug development and molecular diagnostics seem obvious. Unfortunately, these applications of DSC are currently not realistic simply because of the serial nature of sample processing with standard bio calorimeters. i.e. samples can only be processed one at a time per instrument. An array of instruments is logistically impractical. So, realization of the untapped potential of calorimetry in important prescient applications will first require a new generation of DSC instruments. To meet the demands for high-throughput applications these next generation calorimeters must be built with the necessary signal sensitivity and have multiplex capabilities of processing multiple samples simultaneously in parallel.

## Data Availability

The data underlying this article are available in the article and in its online supplementary material.

## Acknowledgements

This work was funded in part by grant No. R43GM146542 from the National Institutes of Health, and an award from the Portland State University Venture Development Fund. We want to thank RayBiotech for supplying insightful samples for our study. The Oregon State University NMR Facility was funded in part by the National Institutes of Health, HEI Grant 1S10OD018518, and by the M.J. Murdock Charitable Trust grant #2014162.

