## Supplemental Material for "Calorimetric analysis using DNA thermal stability to determine protein concentration"

| Table S1: Transition Heat Capacity forced through zero* | | | | |
|---|---|---|---|---|
| Sample | PSV (mL/g) | Slope (µcal mL/mg) | Pearson's r | $C_{p,x}$ (mcal·g$^{-1}$·K$^{-1}$) |
| High Temp DNA Hairpin (NaCl) | 0.55 | 2.378 ± 0.031 | 0.9994 | 2.93 ± 0.03 |
| High Temp DNA Hairpin (CsCl) | 0.55 | 2.358 ± 0.022 | 0.9998 | 2.91 ± 0.02 |
| Low Temp DNA Hairpin (NaCl) | 0.55 | 2.357 ± 0.010 | 0.99997 | 2.91 ± 0.01 |
| Low Temp DNA Hairpin (CsCl) | 0.55 | 2.372 ± 0.037 | 0.9995 | 2.92 ± 0.03 |
| Duplex DNA (NaCl) | 0.625 | 4.095 ± 0.014 | 0.99998 | 4.72 ± 0.01 |
| Human Serum Albumin (NaCl) | 0.733 | 2.122 ± 0.032 | 0.9998 | 2.86 ± 0.03 |
| Human Serum Albumin (CsCl) | 0.733 | 2.140 ± 0.041 | 0.9989 | 2.88 ± 0.04 |
| Lysozyme (NaCl) | 0.703 | 1.921 ± 0.100 | 0.9959 | 2.62 ± 0.10 |

*A line was fit to the $C_p^{max'}$ versus concentration data while being forced to pass through the origin.

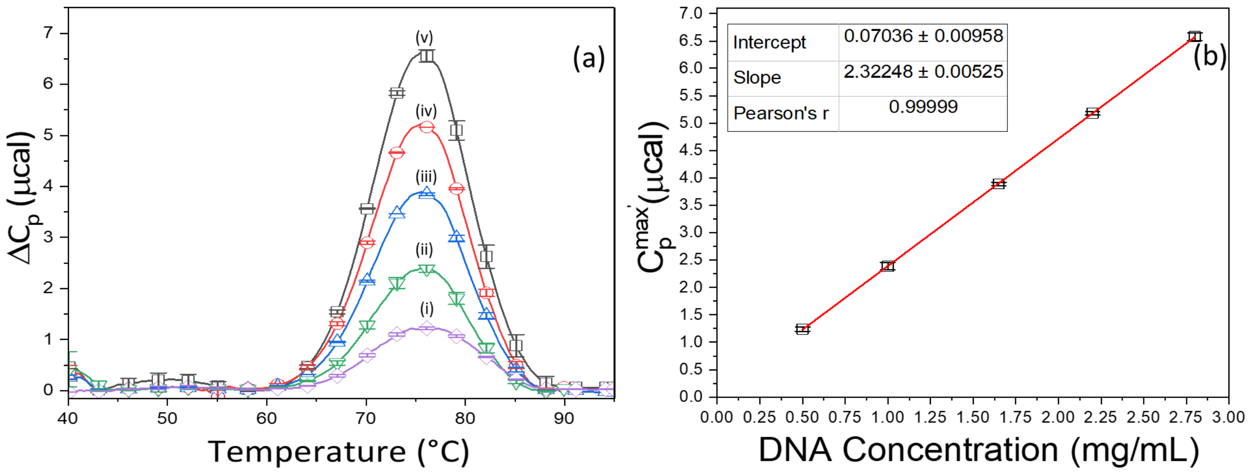


**Figure S1**: (a) Average thermograms for low-temp hairpin concentrations in mg/mL: (i) 0.50, (ii) 1.00, (iii) 1.65, (iv) 2.20, (v) 2.80. (b) Transition heat capacity curve for low-temp hairpin.

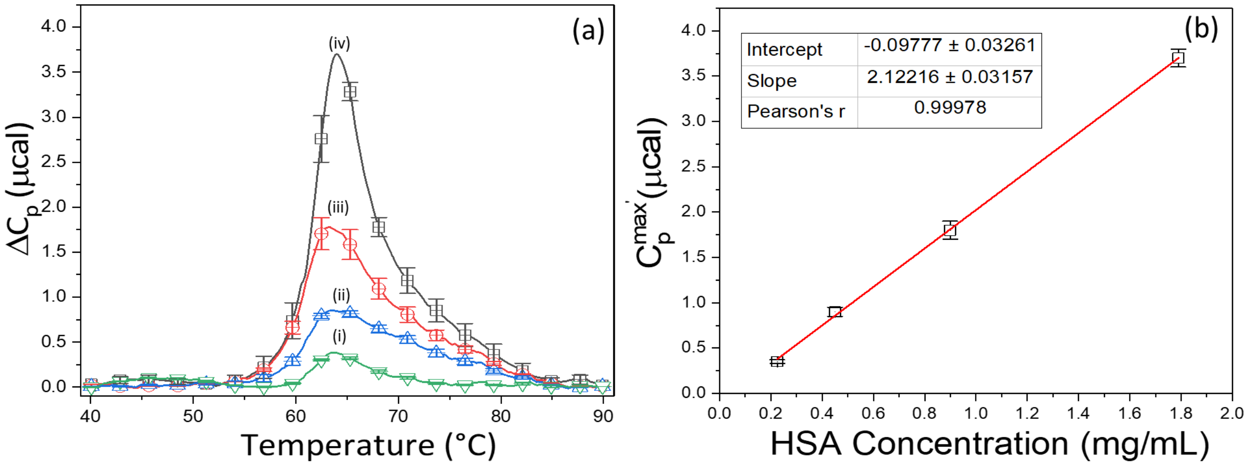


**Figure S2**: (a) Average thermograms for HSA concentrations in mg/mL: (i) 0.225, (ii) 0.45, (iii) 0.90, (iv) 1.80. (b) Transition heat capacity curve for HSA.

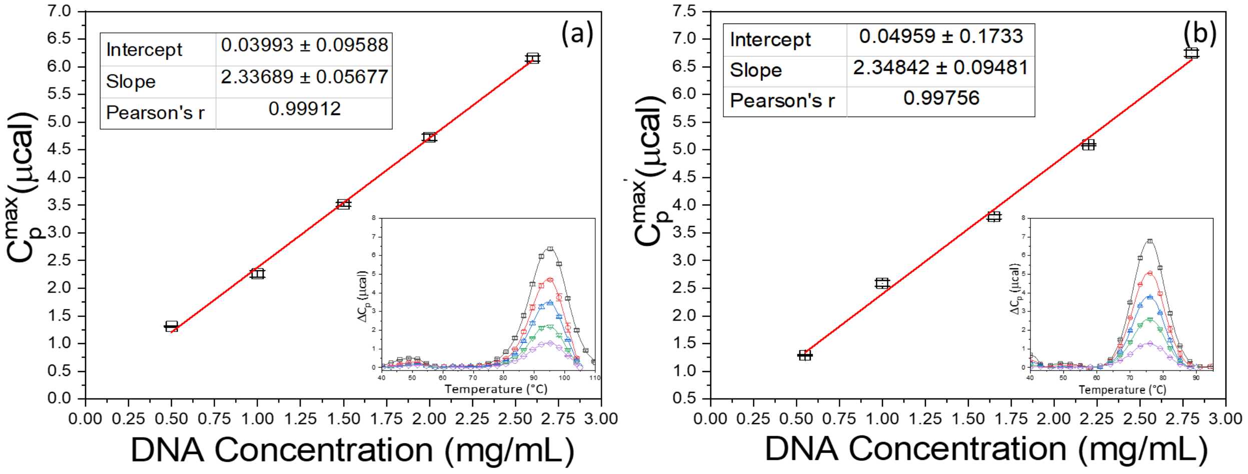


**Figure S3**: (a) Transition heat capacity curve for the high-temp hairpin in CsCl buffer. Inset of (a), the average thermograms for the hi-temp hairpin in CsCl buffer. (b) Transition heat capacity curve for the low-temp hairpin in CsCl buffer. Inset of (b), the average thermograms for low-temp hairpin in CsCl buffer.

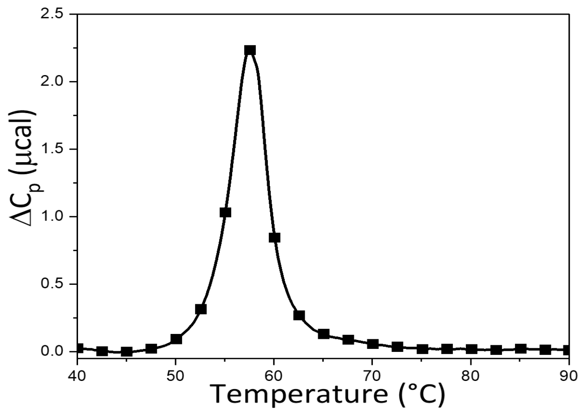


**Figure S4**: Thermogram for 0.40 mg/mL of human herpes simplex virus glycoprotein D (HSV-GpD).
